# Saturation, Task Error, and Feedback Timing Shape Early Implicit Adaptation

**DOI:** 10.1101/2025.09.08.674896

**Authors:** Zacchary Nabaee-Tabriz, Parmin Rahimpoor-Marnani, Alina Khan, Kimerdeep Bassi, B. M. ‘t Hart, Denise Y. P. Henriques

## Abstract

Motor adaptation is essential for maintaining coordination and precision in daily activities. Implicit motor adaptation—adaptation that occurs without conscious awareness—is thought to be primarily driven by sensory prediction errors. Here, we investigated how rapidly these unconscious changes in reaching behavior emerge as a function of error magnitude and the availability of task error signals. To this end, we employed a single-trial learning (STL) paradigm within a classical visuomotor rotation task. Participants made center-out reaching movements to either small (dot) or large (arc) targets while experiencing single perturbation trials with cursor rotations ranging from 1° to 90°, each followed by an aligned washout trial. By manipulating target size, we systematically modulated the presence of task error while holding sensory prediction error constant. We further compared these early implicit changes with those observed during standard prolonged adaptation to a fixed 20° rotation across >100 trials. Our results show that implicit adaptation emerges rapidly, even after a single exposure to small perturbations, and follows a saturating, fixed-rate response profile. Importantly, the magnitude of single-trial adaptation was greater when task error was present (small targets) compared with conditions in which only sensory prediction error was available (large targets). Moreover, STL-derived parameters moderately predicted the initial phase of adaptation during prolonged learning, suggesting that STL captures core dynamics of early implicit processes. These findings provide new insight into the mechanistic principles governing implicit motor adaptation. By identifying the parameters that drive early-stage error-based learning, this work refines current models of sensorimotor learning and highlights potential strategies for designing targeted training or rehabilitation protocols that leverage rapid adaptation processes to enhance motor performance and recovery.

## Introduction

The motor system maintains accurate and coordinated movement by adapting to changes in both the body and the environment. Growth, fatigue, injury, tool use, and changes in environmental context can all alter the mapping between motor commands and sensory consequences, requiring continual recalibration to preserve movement accuracy ((Herzfeld & Shadmehr, 2014; Krakauer et al., 2019)). Motor adaptation supports this flexibility by updating internal models to minimize movement errors and restore precision following altered sensory feedback.

Visuomotor adaptation, in which visual feedback of the hand is perturbed (e.g., by rotating a cursor relative to the true hand position), provides a well-established paradigm for probing these mechanisms. Such perturbations create a mismatch between predicted and actual sensory feedback, initially producing systematic reaching errors. With practice, movements adjust to counteract the perturbation, and residual deviations remain even when the perturbation is removed and participants are instructed to reach directly to the target. These aftereffects reflect implicit changes in motor commands that occur independently of explicit strategy use (Gastrock et al., 2020; Modchalingam et al., 2019, Modchalingam et al., 2023; ‘t Hart et al., 2024; Taylor et al., 2014; Vachon et al., 2020). After prolonged training, aftereffects typically plateau at ∼15° for rotations ≥30° (Bansal et al., 2023; Bond & Taylor, 2015; Modchalingam et al., 2019, Modchalingam et al., 2023; ‘t Hart et al., 2024; Wijeyaratnam et al., 2022). Because they are usually measured only after extensive practice, the time course by which these implicit changes emerge remains unclear, contributing to the view that implicit learning develops slowly and that early adaptation is dominated by explicit strategies (Benson et al., 2011; Bond & Taylor, 2015; Haith & Krakauer, 2014; McDougle et al., 2015). Recent evidence challenges this view, showing that substantial aftereffects can appear within a handful of trials, and may saturate after only one to three exposures (D’Amario et al., 2024; Ruttle et al., 2016, 2021).

Beyond aftereffects, implicit learning has also been probed using error-clamp paradigms, in which the cursor follows a fixed trajectory toward the target regardless of the actual hand movement. Even when explicitly informed that the feedback is uncontrollable, participants gradually shift their unseen hand movements away from the clamped feedback (Al-Fawakhiri et al., 2023; H. E. Kim et al., 2018, 2019; Morehead et al., 2017; Tsay, Haith, et al., 2022; Zhang et al., 2024). These adjustments demonstrate an automatic response to sensory prediction errors—the mismatch between expected and actual sensory outcomes—even in the absence of task-related success or failure, highlighting sensory prediction error as a primary driver of implicit adaptation. However, accumulating evidence indicates that task error—the deviation between the intended movement goal and the actual outcome—can also modulate implicit learning.

Manipulations that minimize task error in classic movement-contingent visuomotor adaptation, such as shifting targets mid-movement or enlarging target size, reliably reduce aftereffects even when sensory prediction error is held constant (H. E. Kim et al., 2019; Leow et al., 2018, Leow et al., 2020, Leow et al., 2025; Tsay, Haith, et al., 2022). This body of work suggests that implicit adaptation is not solely driven by sensory prediction errors, but can be attenuated when task error signals are reduced or eliminated.

To further dissect the contribution of task error, we employed a visuomotor reaching paradigm using single-trial learning (STL). STL measures immediate aftereffects following individual training trials with varying rotation magnitudes, enabling fine-grained tracking of early implicit changes. Critically, we manipulated target size to vary task error: small targets induced task failure when the rotated cursor missed, whereas wide targets minimized task error by ensuring the likelihood of success. This design allowed us to isolate the influence of task error on the earliest stage of implicit adaptation. In addition, by systematically varying rotation magnitude, we examined how the time course of these implicit changes scales with error size, providing insight into the dynamics and limits of trial-by-trial implicit learning.

In addition to error size and task error, the temporal availability of visual feedback may also constrain early implicit adaptation. Delaying feedback has been reported to attenuate aftereffects while leaving explicit strategies intact (Brudner et al., 2016; McDougle & Taylor, 2019; Schween & Hegele, 2017), suggesting that continuous, time-locked error information is critical for implicit recalibration. However, closer inspection of these studies indicates that implicit contributions are not eliminated under delayed feedback, but instead persist at a reduced level. What remains unclear is whether such effects are detectable within a single trial of exposure, and how they interact with error magnitude. In our second experiment, Terminal Feedback Delay, we addressed this question by replacing continuous cursor feedback with terminal endpoint feedback, presented 0–1.6 s after movement completion, in a within-subjects design. Using the STL approach, we tested whether removing online feedback suppresses sensitivity to error magnitude in early implicit adaptation, and whether increasing delay within the terminal range further modulates the size of the implicit update.

Together, these two experiments provide a complementary view of the constraints on early implicit adaptation, revealing how both the nature of the perturbation size, error signal and its temporal availability shape the earliest stages of motor recalibration.

## Methods

### Participants

A total of 179 right-handed undergraduate students from York University participated across two experiments. In Experiment 1: Error Signal, 129 participants (M = 19.79 years, SD = 2.37; 92 female, 36 male, 1 other/prefer not to say) were recruited through the Undergraduate Research Participant Pool (URPP) and the Kinesiology Undergraduate Research Experience (KURE). In Experiment 2: Terminal Feedback Delay, 50 participants (mean age ≈ 20 years) were recruited through the URPP. All participants reported normal or corrected-to-normal vision and provided informed consent prior to participation. The study protocol was approved by the York University Human Participants Research Ethics Board.

### Apparatus

Both experiments used the same setup. Participants performed reaching movements with a handheld stylus on a Wacom Intuos Pro Large digitizing tablet (44 × 27 cm) as shown in Fig 1B. Visual stimuli were displayed on a downward-facing monitor (DELL E2009W, 44.3 × 24.9 cm) positioned 25 cm above the tablet, and reflected onto a mirror placed midway between the monitor and tablet. This arrangement made the stimuli appear in the plane of the hand while occluding direct vision of the arm.

**Figure 1.**
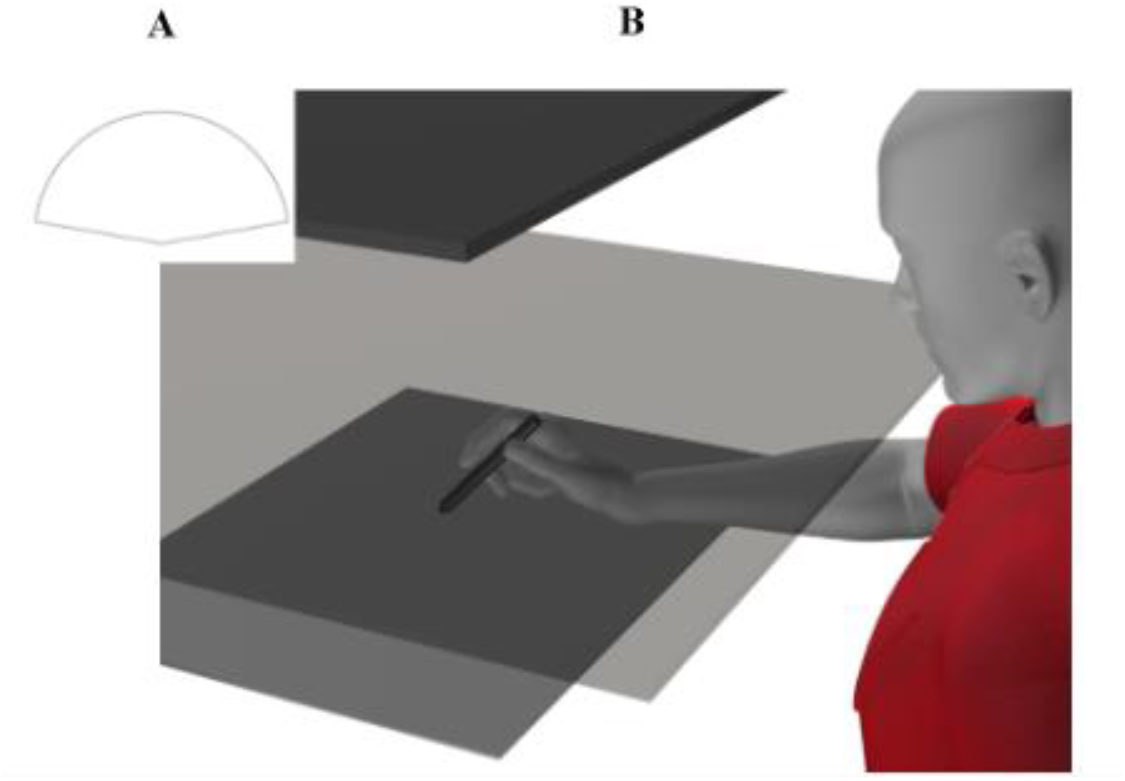
Apparatus used for the experiment. A: Cone-shaped stencil placed on the digitizing tablet to constrain movement amplitude and guide the hand back to the center. B: The stylus was moved along the digitizing tablet while the cursor position, projected from the monitor above, was reflected off a mirror that occluded the participant’s hand.

A plastic stencil with a cone-shaped opening (8.5 cm radius, spanning 160°, Fig 1A and 2) constrained movement amplitude and guided return to a fixed home position at the center of the workspace. Targets were presented 8 cm from the home position at one of five angular locations (60°, 75°, 90°, 105°, or 120°). The cursor was a 3 mm radius disk; the home and target circles were outlined but not filled.

## Experimental Procedure

Participants made rapid, straight reaches to targets using a stylus-controlled cursor. The start position was indicated by a blue circle (5 mm radius), and targets appeared either as a small dot (3 mm radius) at one of five angles (60°, 75°, 90°, 105°, or 120°) for both Experiments as shown in Fig 2A or as a wide arc spanning 120° centered at 90° as shown in Fig 2B for part of Error-signal Experiment.

**Figure 2.**
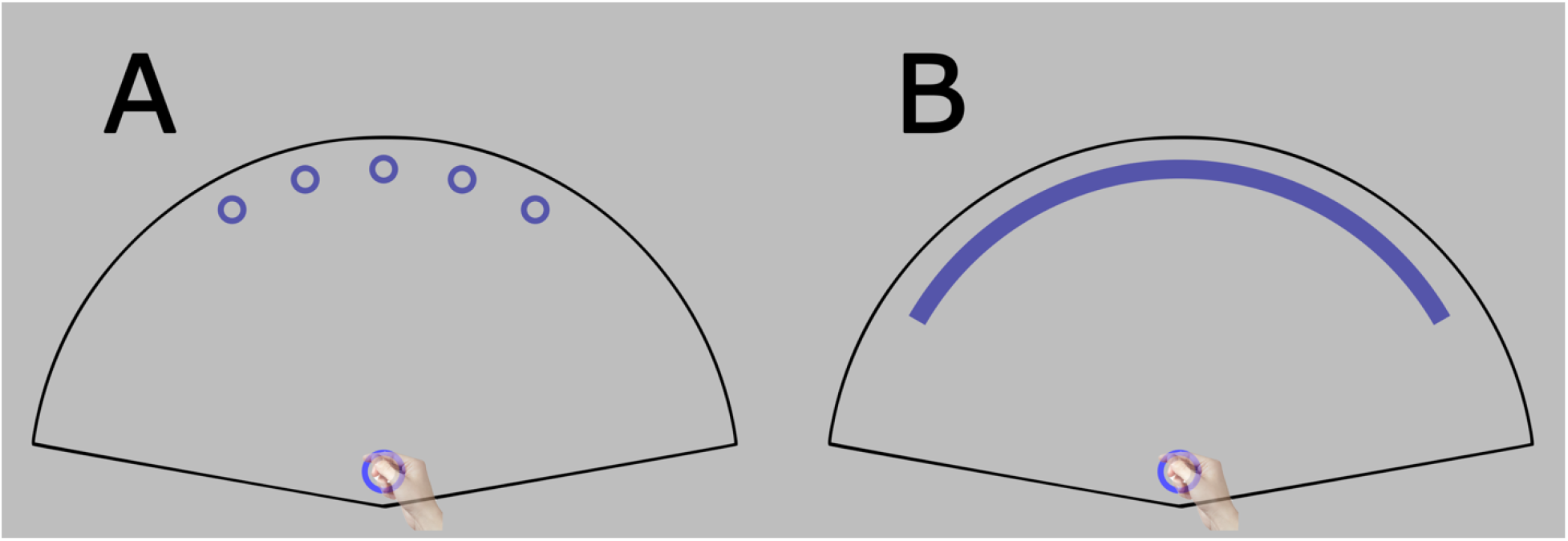
**(A)** Visual example of all possible dot target locations (60º, 75º, 90º, 105º, or 120º), followed by a visual example of the arc target **(B)**. Any part of the arc target may be reached. The 5 mm blue circle represents the home position, and the white circle represents the cursor. All targets are located 8 cm away from the home position. The stencil, which serves as the area where reaches may be done, surrounds both the dot and arc targets, as well as the home position. An added advantage of this cone-shaped stencil design is that it guides the stylus back to the home position following a reach, even when the cursor is not visible.

Each trial began when the participant held the stylus at the home position for 0.1 s. When the target appeared, participants initiated a reach. Once the stylus traveled 0.5 cm radially, the home position disappeared, and the trial ended when the stylus crossed the target radius (8 cm). After trial completion, the target and cursor imprint remained visible for 0.6 s before the participant returned to the home position to begin the next trial.

Four trial types were used: baseline trials with veridical feedback to assess unperturbed reaching, rotation trials introducing a visuomotor perturbation, aftereffect trials without feedback to measure implicit adaptation, and washout trials with veridical feedback to reduce residual learning. These trial types were combined into short single-trial learning (STL) bouts of 4–6 trials: a baseline trial, a rotation trial with a pseudorandomly selected magnitude, an aftereffect trial, and 1–3 washout trials. Rotation trials were directed to a single dot or arc target centered at 90°, whereas surrounding unperturbed trials were directed to one of five dot targets. The number of washout trials, as well as the rotation magnitudes and directions (clockwise or counterclockwise), were pseudorandomized. Each magnitude was presented five times per direction for both dot and arc targets.

Both experiments consisted of three phases (see Fig. 3 for an example schedule from the 60°-max group). In the familiarization phase (80 trials, prior to the vertical dashed line in Fig. 3), participants practiced reaching with veridical feedback, first to dot targets (40 trials) and then to arc targets (40 trials), to acclimate to the setup. The STL phase (∼1200 trials) then followed, comprising repeated STL bouts with pseudorandomized rotation magnitudes and directions. Target type alternated between bouts, indicated by line color in Fig. 3 (blue for dot targets, orange for arc targets).

**Figure 3.**
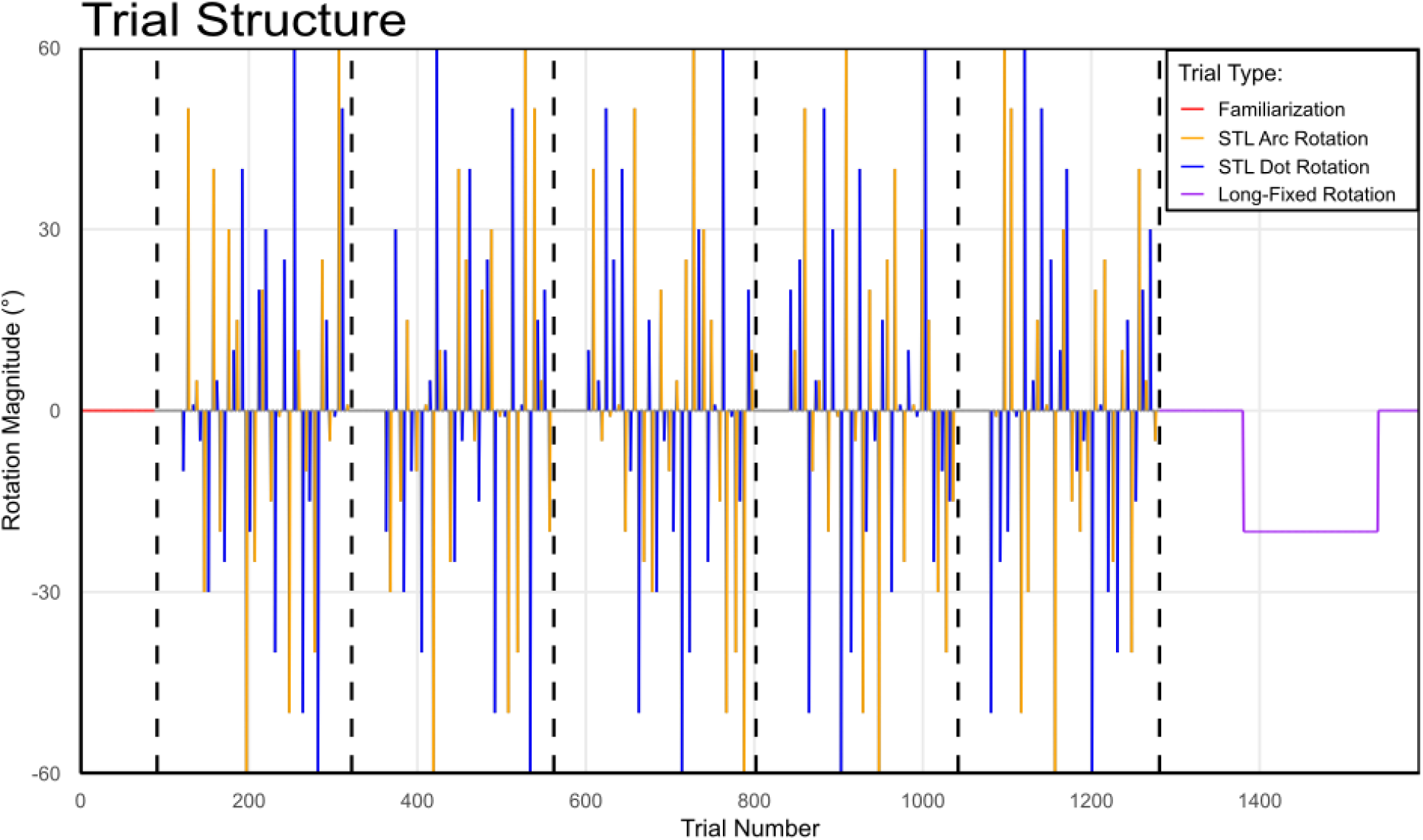
An example of the perturbation schedule from the three main experimental phases. The Familiarization phase **(trials 0 – 80)**, consists of 40 familiarization trials with the circular target, followed another 40 trials with the arc target. The STL phase **(trials 81 – 1281)**, consists of 1200 trials. STL bouts are made up of 1 baseline, 1 rotation, 1 aftereffects, and 1 – 3 washout trials. The rotation values are pseudorandomized from bout to bout. The Long-Fixed Rotation Phase **(trials 1282 – 1591)** consists of 310 trials: 40 aligned-acquire trials, 60 baseline trials, 160 rotation trials with a fixed rotation of -20°, followed by 25 after-effects and 25 washout trials. Black vertically dashed lines represent mandatory 1-minute breaks. The first 40 trials following each break consisted of 10 aligned-acquire trials, followed by 30 baseline trials.

In the Error Signal experiment, a fourth phase was included to assess whether performance during STL bouts generalized to prolonged exposure. This long-fixed rotation (LFR) phase (310 trials, right side of Fig 3) consisted of 40 aligned-acquire trials, 60 baseline trials, 160 rotation trials with a fixed 20° rotation, followed by 25 aftereffect and 25 washout trials.

To minimize fatigue in both experiments, mandatory one-minute breaks were introduced approximately every 240 trials during the STL phase (indicated by vertical dashed lines in Fig. 3). After each break, participants completed a short block of baseline reaches in which the cursor was required to acquire the target, reinforcing accurate movements before resuming STL bouts or continuing the LFR phase.

In the Error signal experiment, participants were assigned to one of three rotation-magnitude groups:

45°-max group: ±1°, 5°, 10°, 15°, 20°, 25°, 30°, 35°, 40°, 45° (n = 30)

60°-max group: ±1°, 5°, 10°, 15°, 20°, 25°, 30°, 40°, 50°, 60° (n = 43)

90°-max group: ±1°, 5°, 10°, 15°, 20°, 30°, 40°, 50°, 70°, 90° (n = 56)

resuming the subsequent set of STL bouts and prolonged training task.

In the Error Signal experiment, in rotation trials, dot and arc targets were used to manipulate task error. Because rotation magnitude and direction were randomized, reaches to dot targets almost always missed, producing task error (Fig. 4A). In contrast, the wide arc target ensured a high likelihood of cursor–target overlap even for large rotations, minimizing task error (Fig. 4B). Importantly, sensory prediction error was present for both target types due to the cursor rotation. Implicit adaptation to a single rotation was quantified as the change in hand angle between the baseline trial preceding the rotation and the aftereffect trial that followed it. Comparing these single-trial aftereffects following rotated reaches to dot and arc targets assessed the contribution of task error to initial implicit adaptation, while analyzing them across rotation magnitudes allowed us to examine how error size influences the time course and magnitude of adaptation.

**Figure 4.**
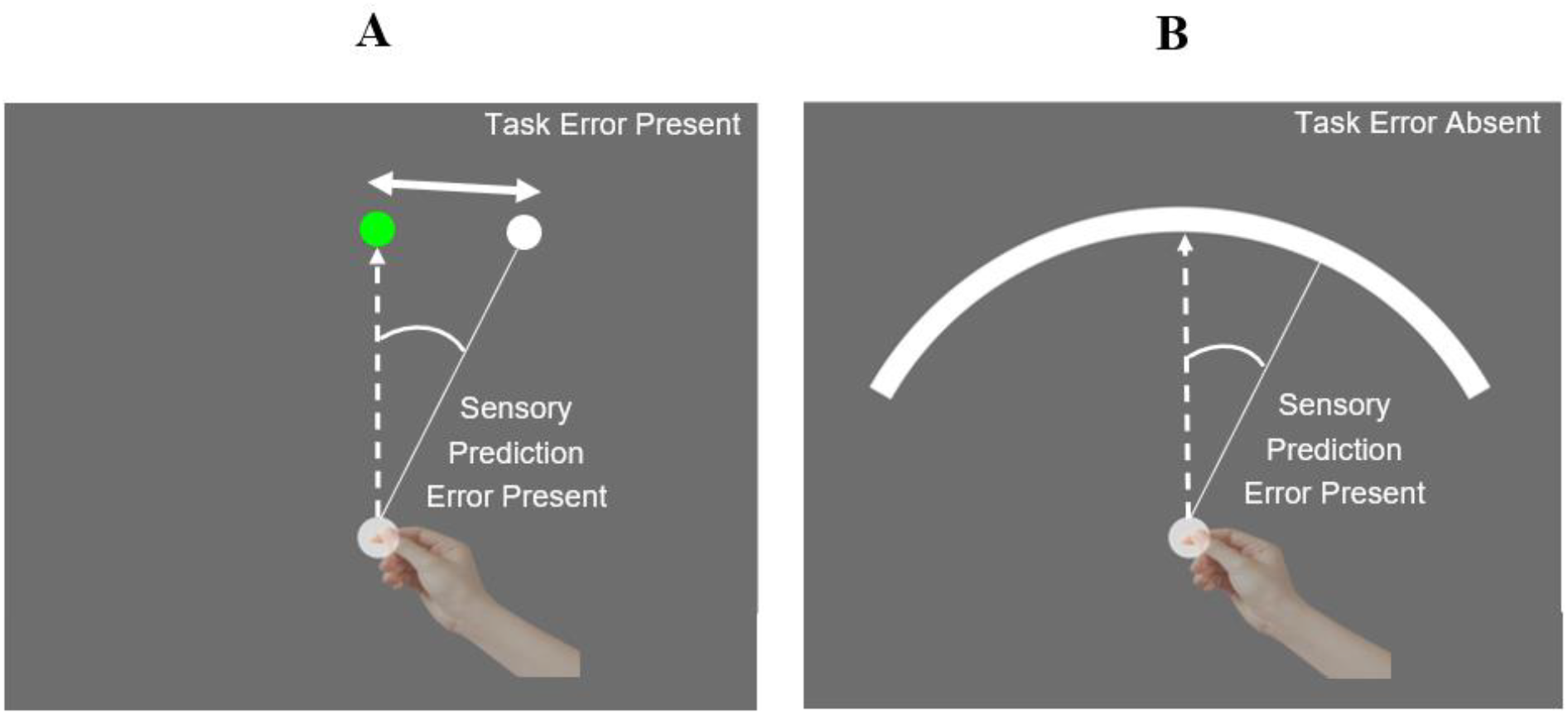
Visual depiction of how both sensory prediction error and task error are present when reaching towards circular targets on Rotation trials (A), whereas the arc targets (B) greatly reduce the likelihood of task error by providing a broad region for successful hits. The hand used for making reaches is occluded throughout the duration of the experiment. The dashed line represents hand trajectory and the solid line represents cursor’s trajectory.

In the Terminal Feedback Delay experiment, participants completed the same three phases and rotation magnitudes as the 45°-max group of the Error Signal experiment, which served as the continuous-feedback control. For the terminal feedback group, outward reaches were made without visual feedback until the stylus contacted the stencil. At that point, the target disappeared and the pink feedback cursor appeared at the endpoint location only after a randomized delay (0.0, 0.2, 0.4, 0.8, or 1.6 s). The cursor then vanished, and participants returned to the home position to begin the next trial. Thus, the task was identical to Experiment 1 except that on rotation trials feedback was terminal, delayed by one of five intervals. All STL trials in this experiment used dot targets, with 240 STL bouts, for a total of 1,380 trials including baseline and washout trials).

### Data Analysis

Reach direction was computed online when the cursor had moved 2 cm from the start position, approximately one-quarter of the total 8 cm movement distance. This point precedes peak velocity and minimizes the influence of late trajectory corrections. Angular deviation was calculated relative to the target direction. Initial implicit change for each bout was obtained by subtracting the baseline deviation from the corresponding aftereffect deviation. Baseline and aftereffect trials were identical in terms of target type (dot target) and movement criteria, allowing a direct estimate of the implicit change produced by a single rotation exposure.

Rotation magnitudes for all STL bouts within a participant’s session were extracted for analysis. Baseline and aftereffect trials with deviations exceeding ±45° were excluded as outliers. For visualization and analysis, data were collapsed across clockwise and counterclockwise rotations, normalized to a positive scale, and averaged by rotation magnitude. All statistical analyses were performed in R (https://www.r-project.org/).

#### Effect of error-signal type on initial aftereffects

To examine the effect of error-signal type in Experiment 1, separate two-way repeated-measures ANOVAs were conducted for each rotation-magnitude group (45°-max, 60°-max, and 90°-max), with target type (dot, arc) and rotation magnitude (10 levels per group) as within-subject factors. Rotation-magnitude groups were analyzed independently and not directly compared.

#### Modeling the effect of rotation size on initial adaptation

To examine how initial implicit adaptation varies with rotation size, we fitted each participant’s STL aftereffect data to two competing models reflecting previously observed attenuation and saturation of learning with increasing error: an attenuation model and a capped fixed-rate model (Fig. 5).

**Figure 5.**
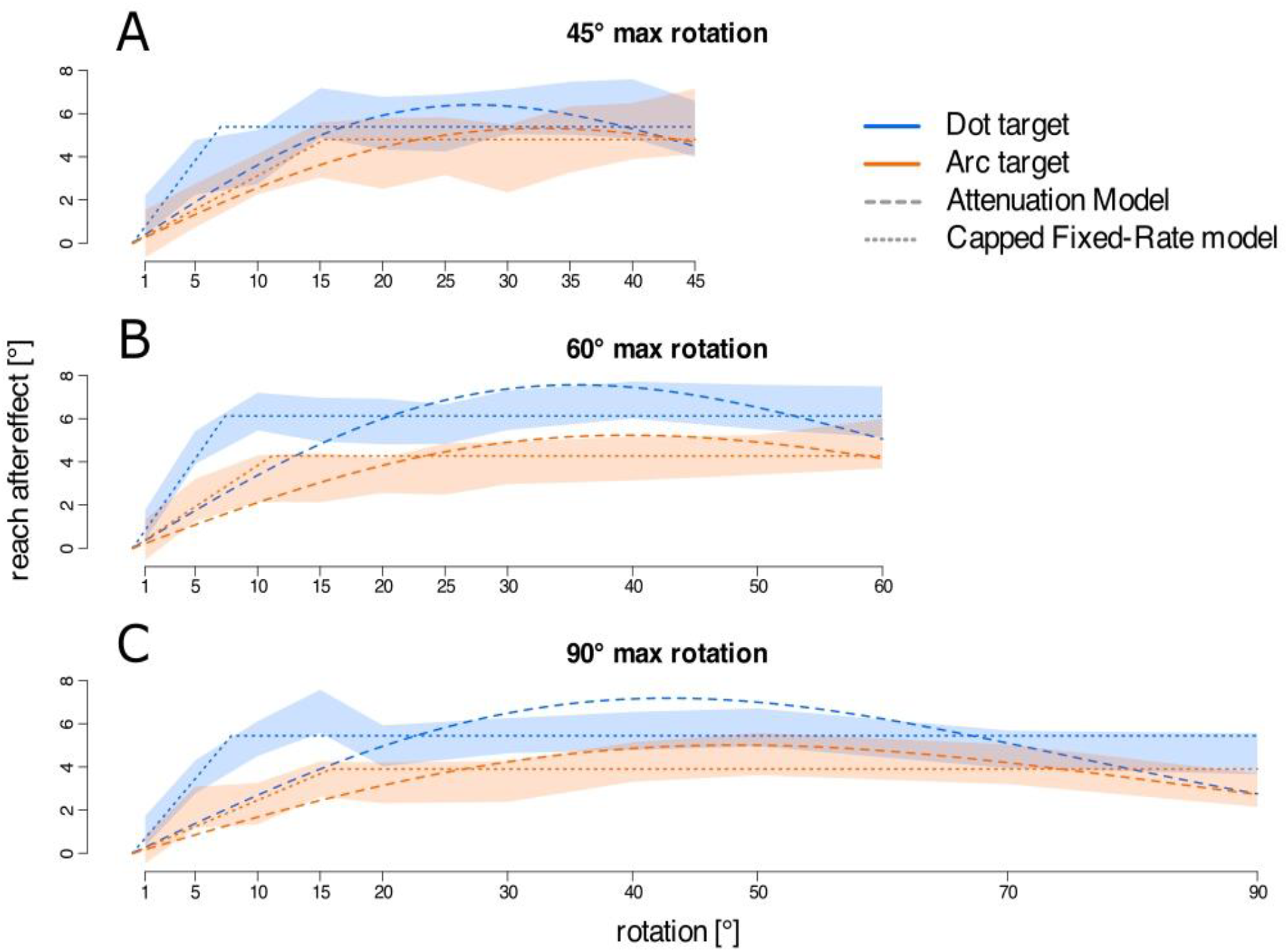
Averaged aftereffects and fitted model predictions of the capped fixed-rate (dotted lines) and attenuation (dashed lines) models for the 45° group **(A)**, the 60° group **(B)**, and the 90° group **(C)**. Plotted separately for dot (blue) and arc (orange) targets. Shaded regions indicate the bootstrapped 95% confidence interval of the group means.

The attenuation model assumes that very large errors are less likely to be attributed internally, resulting in reduced implicit adaptation. It is expressed as:

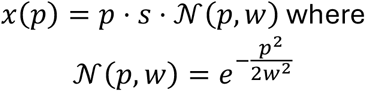

Here, *x(p)* is the predicted reach deviation for a perturbation of size ppp, sss is a scaling factor representing the fraction of error corrected when attributed internally, and 𝒩 reflects error sensitivity (variance of the Gaussian density function). Normalization ensures the function equals 1 at *p*=0, meaning small rotations are corrected proportionally but implicit learning attenuates for larger errors.

The capped fixed-rate model assumes that implicit adaptation increases linearly with error up to a fixed maximum, beyond which it saturates:

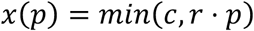

where *c* is the cap level, *r* is the fractional correction rate, and *p* is the perturbation size.

For each participant, parameter sets (c, r) for the capped model and (s, w) for the attenuation model were estimated using grid search followed by bounded optimization. The 10 parameter sets yielding the lowest mean squared error (MSE) were refined to obtain a single best-fitting parameter set per model and participant. Group-level model fits were compared using MSE and corrected Akaike Information Criterion (AICc) values, and paired-sample t-tests on individual MSE values assessed relative goodness of fit.

#### Predicting prolonged adaptation from STL

We next tested whether STL-based model predictions generalize to adaptation during prolonged exposure. During the long-fixed rotation (LFR) phase, participants completed 160 trials with a constant 20° perturbation. For each participant, adaptation after the first rotation trial was estimated by fitting an exponential function to their full learning curve and extracting the predicted value after the first perturbation.

Similarly, for STL data with dot targets only, both models were fitted to each participant’s single-trial aftereffects, and the fitted value for a 20° rotation was extracted. If STL reliably predicts prolonged learning, these two estimates should match, falling along a unity line (slope = 1, intercept = 0). To evaluate this, we regressed the LFR exponential estimates onto the STL model predictions (intercept constrained to 0) for both models separately. The slope quantified predictive accuracy: a slope of 1 indicates perfect prediction, slopes <1 indicate overestimation by STL models, and slopes >1 indicate underestimation. Model performance in this context was compared using AICc and R^2^ from the regression fits.

Finally, to rule out underestimation from the exponential fits, we directly compared each participant’s exponential estimate to their actual reach deviation on the first rotated trial of the LFR phase. Paired-sample t-tests assessed whether these differed significantly, clarifying whether any observed overprediction by STL models reflected genuine differences in adaptation dynamics or curve-fitting artifacts. All data are available in this repository: https://osf.io/6m24e/

#### Effect of delayed terminal feedback on initial aftereffects

To assess the effects of feedback delay and perturbation size in Experiment 2, a two-way repeated-measures ANOVA was conducted within the terminal feedback group, with feedback delay (0.0, 0.2, 0.4, 0.8, 1.6 s) and rotation magnitude (six levels) as within-subject factors. To evaluate the influence of feedback type, aftereffects were collapsed across delay durations and compared with the continuous-feedback group from Experiment 1 using a mixed-design ANOVA, with feedback type (terminal, continuous) as a between-subject factor and rotation magnitude as a within-subject factor.

### Results

### Error Signal Experiment

#### Effect of error signal and perturbation size on implicit motor adaptation

Significant aftereffects were observed following a single exposure to a cursor rotation (Fig. 5). The relationship between perturbation size and initial aftereffects was broadly similar across the three rotation-range groups, with comparable patterns for both dot (blue curves) and arc targets (orange curves). Across all groups, however, dot-target training produced consistently larger aftereffects than arc-target training—by approximately 1–2°—in the 45° group (Fig. 5A) [F(1,30) = 11.52, p = 0.002, η^2^g = 0.027], the 60° group (Fig. 5B) [F(1,42) = 66.24, p < 0.001, η^2^g = 0.103], and the 90° group (Fig. 5C) [F(1,55) = 19.27, p < 0.001]. Despite this small but reliable main effect of target type, no significant interaction between target type and rotation magnitude was found in any group, indicating that the shape of the aftereffect function was similar across conditions. The main effect of target type suggests that additional task error signals in the dot-target condition yielded slightly larger initial implicit changes than trials involving only sensory prediction error, as in the arc condition.

#### Effect of rotation size on implicit adaptation: capped fixed-rate vs. attenuation model

To assess how initial changes scaled with perturbation size, we fit attenuation and capped fixed-rate models to STL aftereffects. Data from dot and arc conditions were combined, as no interaction with rotation magnitude was observed. Both models captured the overall pattern, but the capped fixed-rate model provided a closer match: adaptation increased with error size before plateauing at ∼5°–6°, whereas the attenuation model predicted a decline at larger rotations not seen in the data (Fig. 6). This was confirmed quantitatively, with lower mean squared error (MSE) and Akaike Information Criterion (AIC) values for the capped fixed-rate model (paired t-test on MSEs: t(257) = –9.12, p < 0.001, 95% CI [–2.10, –1.36]). Participant-level comparisons (Fig. 6) showed most points near the unity line, but a subset with markedly higher prediction error for the attenuation model, again indicating a better fit for the capped fixed-rate model. Together, these results suggest that initial implicit adaptation grows with error size up to a fixed limit, beyond which larger perturbations do not elicit stronger aftereffects.

**Figure 6.**
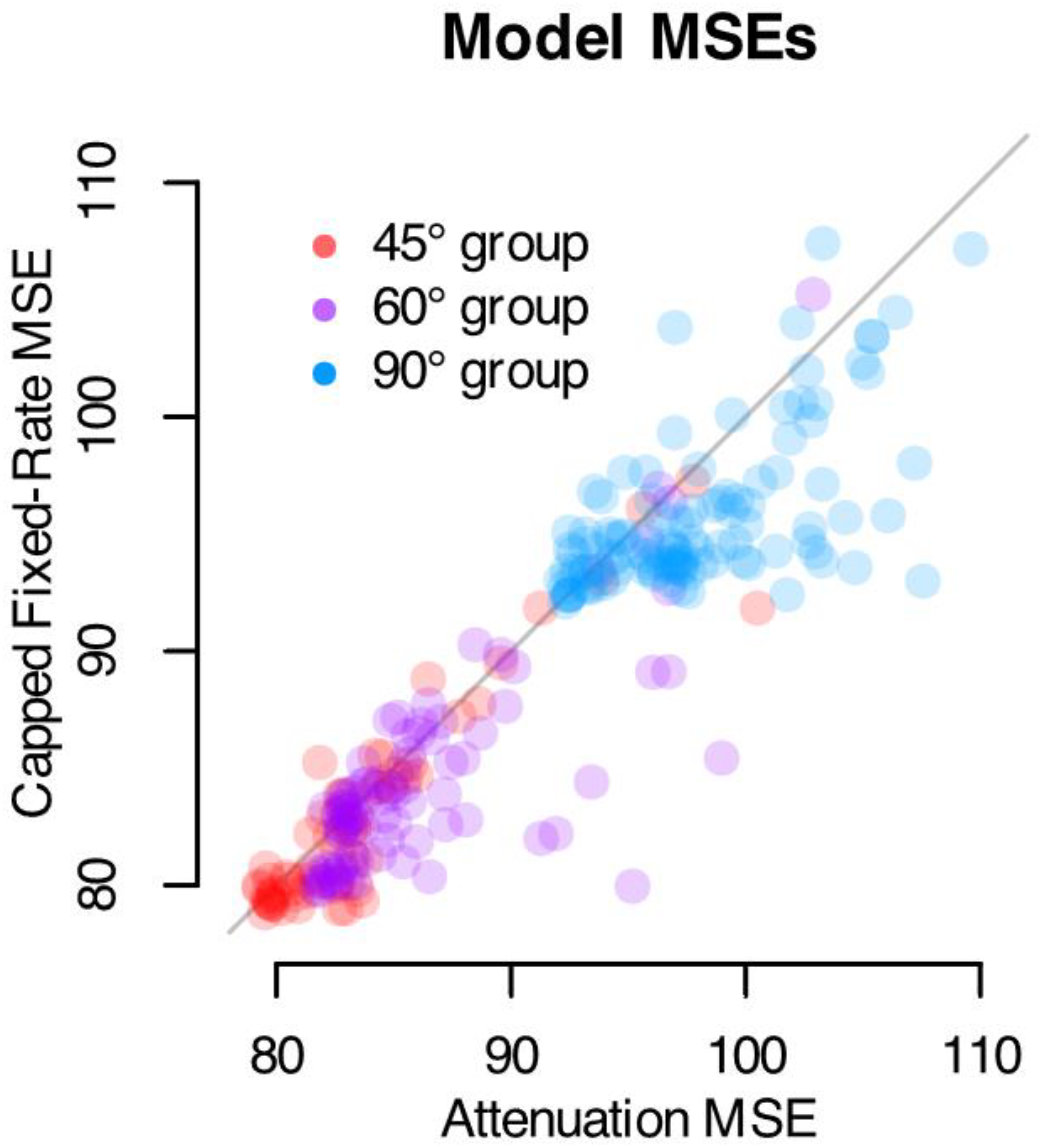
Model fit comparison across participants. Each point represents a single participant’s mean squared error (MSE) for the capped fixed-rate model (y-axis) and the attenuation model (x-axis), computed from single-trial learning fits between the 45° **(Red)**, 60° **(Purple)**, and 90° **(Blue)**, rotation groups. The diagonal line indicates the unity line (i.e., equal model fits). Points located further away from the unity line indicated a poorer fit with respect to the model they are closer to.

We next tested whether STL-derived model fits could predict early adaptation in a long-exposure (LFR) setting. For each rotation group, model predictions based on STL data were compared with estimates of initial adaptation during a 200-trial block with a 20° rotation. Predictions from both models correlated significantly with observed long-exposure adaptation, with slopes ranging from 0.28 to 0.56, although STL consistently overestimated the magnitude. This overestimation was not explained by bias in the exponential decay function used to estimate initial adaptation, as direct comparisons with the first rotated trial showed no significant difference and, if anything, a slight overprediction. These results suggest that STL can capture individual differences in early adaptation and generalize to extended training, though adjustments may be needed to account for systematic overestimation.

#### Predicting Implicit Adaptation in Prolonged Exposure Training from Single-Trial Learning

We tested whether single-trial learning (STL) predicts adaptation in a 160-trial fixed-rotation block with a 20° visuomotor rotation. Early adaptation from prolonged training was estimated for each participant by fitting an exponential decay function to the learning curve and taking its value after the first rotated trial. Predictions, based on dot-target trials only, were generated for each rotation-size group.

As shown in Figure 7 (left column), adaptation to the 20° rotation was robust across groups. Slopes relating STL predictions to prolonged-training estimates were significantly positive for both the capped fixed-rate and attenuation models (all p <.001): 45°-max group: 0.32 vs. 0.28; 60°-max: 0.47 vs. 0.46; 90°-max: 0.56 vs. 0.53. R^2^ values ranged from.39–.59, with slightly better fits for the capped fixed-rate model (lower AICs in two of three groups).

**Figure 7.**
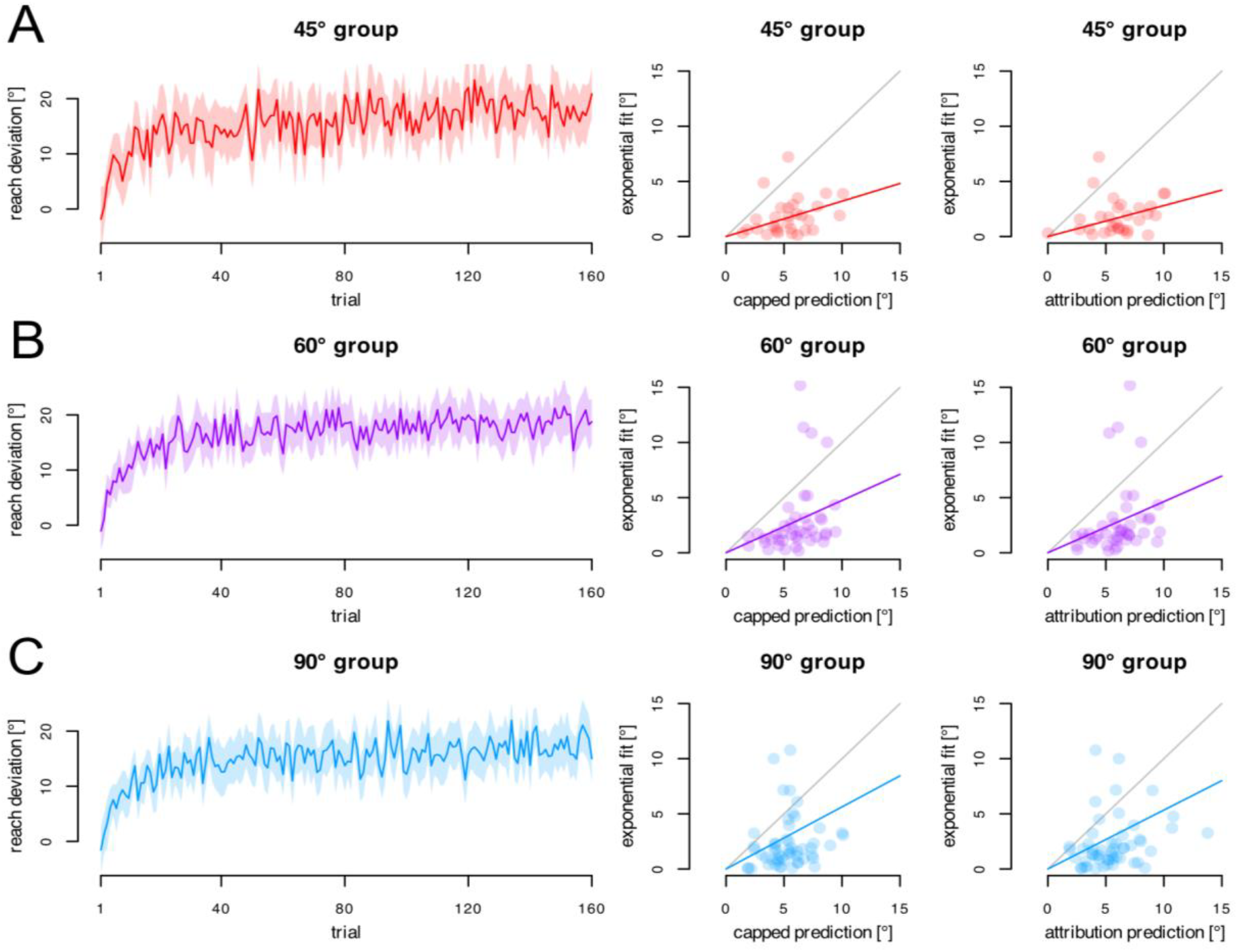
Predicted initial adaptation in the long-fixed rotation phase based on single-trial learning across the across the 45° (A), 60° (B), and 90° (C) rotation groups. Left panels depict average reach deviations over 200 trials in the long exposure block, with shaded regions representing 95% CI and superimposed exponential fits. Middle and right panels show the predicted reach deviations from the capped fixed-rate and attenuation models, respectively, plotted against corresponding exponential fit estimates.

Although STL predictions correlated strongly with early adaptation, all slopes were well below 1.0, indicating overestimation of adaptation in prolonged training. This pattern is also evident in Figure 7 (middle and right columns), where many points fall below the identity line.

### Terminal-feedback delay Experiment

As shown in Figure 8 (purple curves), aftereffects under the terminal feedback condition were small—on the order of a few degrees—but the confidence intervals lay entirely above zero, indicating a statistically significant implicit adaptation effect. To assess whether feedback delay modulated these aftereffects across different rotation sizes, we conducted a two-way repeated-measures ANOVA with within-subject factors of feedback delay and rotation magnitude, applying Greenhouse–Geisser corrections where appropriate. Contrary to our expectation that longer feedback delays would progressively impair these immediate implicit adaptation, the aftereffect functions were highly similar across delay durations (Fig. 8). The main effect of delay duration was not significant, F(3.43, 168.02) = 1.60, p = 0.185, ges = 0.005, indicating that once visual feedback was restricted to the terminal period, the exact timing of its presentation did not further influence implicit learning.

**Fig. 8.**
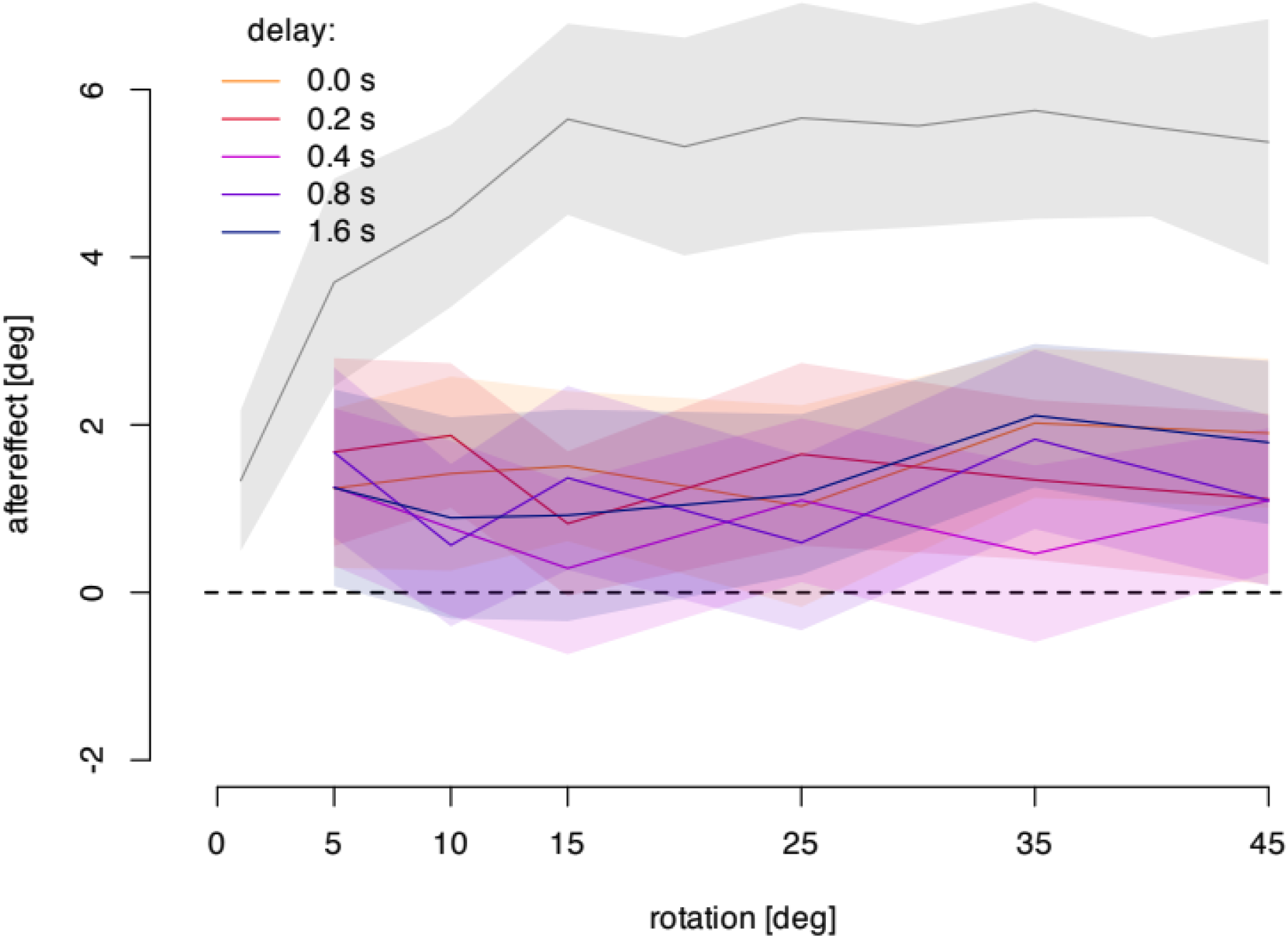
Mean aftereffects (°) plotted as a function of rotation magnitude (5°, 10°, 15°, 25°, 35°, 45°), separately for each of the five terminal feedback delay groups (0.0, 0.2, 0.4, 0.8, 1.6 s). The gray curve represents a control group that received continuous cursor feedback throughout the movement. Terminal feedback resulted in reduced aftereffects across all rotation magnitudes compared to continuous feedback. Importantly, no systematic differences were observed across the five delay durations, indicating that increasing delay alone did not modulate implicit adaptation. Shaded regions indicate the bootstrapped 95% confidence interval of the group means.

Aftereffects within the terminal feedback condition also did not vary systematically with rotation size, as reflected in the relatively flat functions in Figure 8. This was confirmed by a non-significant main effect of rotation magnitude, F(4.25, 208.41) = 1.09, p = 0.362, ges = 0.003, and a non-significant delay × rotation interaction, F(13.50, 661.58) = 0.74, p = 0.730, ges = 0.009. These results suggest that implicit adaptation under terminal feedback was uniformly reduced across all feedback delays and did not scale with error size.

#### Comparison of Terminal and Continuous Feedback

Because feedback delay had no significant effect within the terminal feedback condition, aftereffects were averaged across all delay durations for comparison with an independent group that received continuous visual feedback throughout the movement. A two-way mixed-design ANOVA with feedback group (terminal vs. continuous) as a between-subjects factor and rotation magnitude as a within-subjects factor revealed a robust main effect of group, F(1, 79) = 149.09, p < 0.001, ges = 0.359, confirming that aftereffects were substantially smaller with terminal feedback. This effect size indicates that feedback type accounted for ∼36% of the variance in aftereffects.

There was also a main effect of rotation magnitude, F(4.27, 337.09) = 2.43, p = 0.044, and a significant group × rotation interaction, F(4.27, 337.09) = 2.81, p = 0.023. Continuous feedback produced larger aftereffects that scaled modestly with rotation size, whereas terminal feedback flattened this scaling relationship, eliminating the typical increase in adaptation with larger errors.

## Discussion

Using a single-trial learning (STL) paradigm, we investigated how early implicit adaptation is shaped by the size of sensory prediction errors, the presence of task errors (Error Signal experiment), and the timing of visual feedback (Feedback Delay experiment). By interleaving rotated-cursor trials with veridical feedback trials, we isolated initial reach aftereffects across a broad range of visuomotor perturbations under conditions that either permitted or minimized task error. The Error Signal experiment demonstrated that adaptation increases with error size before saturating, and that task error provides a consistent, additive boost from the very first trials. The Feedback Delay experiment revealed that terminal feedback yields smaller but reliable aftereffects compared with continuous feedback, and that these aftereffects remain robust even when feedback is delayed by up to 1.2 seconds. Together, these findings establish that early implicit adaptation is both bounded by error magnitude and resilient to substantial delays in visual feedback, while being strongly facilitated by the presence of task error.

In the Error Signal experiment, aftereffects increased for small perturbations before plateauing at ∼6°, a pattern captured more accurately by a capped fixed-rate model than by an attenuation model that predicts declines at large errors. Significant adaptation occurred even for the smallest perturbation (1°) with dot targets, indicating that the motor system adjusts to minute discrepancies after a single exposure. This sublinear error–adaptation function mirrors results from long-exposure and error-clamp paradigms (Bond & Taylor, 2015; D’Amario et al., 2024; Hutter & Taylor, 2018; Modchalingam et al., 2019, Modchalingam et al., 2023; ‘t Hart et al., 2024; Werner et al., 2015), but our findings, like those mentioned below, demonstrate that it emerges immediately, without the need for extended training. The magnitude of these immediate aftereffects aligns with other STL studies—1–2° for rotations under 10° (Esfandiari et al., 2022) and ∼5° for rotations of 10–15° (O. A. Kim et al., 2022; Vandevoorde & Orban de Xivry, 2019) —but is nearly tenfold larger than the values reported with STL of clamped rotations ranging from 3.5° to 90° (Tsay et al., 2021).

Importantly, reducing task error by ensuring target–cursor contact (arc targets) consistently reduced aftereffects relative to conditions in which misses were possible (dot targets), despite identical sensory prediction errors. This fixed 1–2° boost (10–20% greater adaptation) across rotation sizes suggests that task error contributes additively rather than scaling with error magnitude. Taken together, these results highlight two core features of early implicit adaptation: a rapid saturation limit evident on first exposure, and an independent contribution of task error that enhances recalibration beyond prediction error.

This early plateau aligns with the view that implicit recalibration is inherently constrained, likely by limits on how the system encodes or utilizes sensory prediction errors once they exceed a critical magnitude. Previous studies using prolonged exposure have reported saturation at higher values (∼20°; (Bond & Taylor, 2015; H. E. Kim et al., 2018; Modchalingam et al., 2019, Modchalingam et al., 2023; Morehead et al., 2017; Tsay, Haith, et al., 2022), while others have argued for attenuation at large errors due to external error attribution (Körding et al., 2007; Tsay et al., 2021; Wei & Körding, 2009; Zhang et al., 2024). Our STL results suggest instead that saturation is a fundamental property of the implicit system itself, evident from the very first trial, and not the cumulative result of extended practice or attributional processes. Furthermore, model fits to STL data predicted adaptation during long-exposure training, indicating that STL provides a rapid and efficient method for mapping adaptation profiles, with potential utility for both mechanistic studies and clinical assessment.

The Feedback Delay experiment complements this perspective by showing that implicit adaptation is highly sensitive to the temporal availability of feedback, but only up to a point. Consistent with earlier work, terminal feedback substantially reduced adaptation compared to continuous feedback. However, once feedback was terminal, increasing the delay up to 1.2 seconds had no additional effect: aftereffects were suppressed but stable across all delays and perturbation sizes. This plateauing effect clarifies the temporal dynamics of implicit recalibration, suggesting that the loss of online cursor information imposes a categorical limit beyond which further delays are functionally irrelevant.

Notably, our results conflict with prior claims that delayed visual feedback eliminates or nearly abolishes implicit learning (Brudner et al., 2016; McDougle & Taylor, 2019; Schween & Hegele, 2017; Tsay, Schuck, et al., 2022). Closer inspection, however, shows that each of these studies actually reported significant aftereffects under delayed terminal feedback, though typically smaller or more transien. For example, Schween and Hegele (2017) found that aftereffects decreased from 11.1° with continuous feedback to 7.2° and 4.0° with 0.2-s and 1.5-s delayst. Brudner et al. (2016) likewise reported immediate aftereffects of ∼10° under both continuous and delayed terminal feedback, with faster decay during washout in the latter. McDougle and Taylor (2019) also observed modest, aftereffects (∼2.7°) after 2-s delays, though their values may underestimate the true magnitude due to averaging across many washout trials. Even Tsay et al. (2022), despite concluding otherwise, showed plots where aftereffects in controls remained above baseline. Consistent with this reinterpretation, Avraham et al. (2020) found that the initial direction of unseen reaches was unaffected when terminal feedback was delayed up to 2 s, suggesting implicit adjustments are preserved. A recent study from our lab (D’Amario et al., 2024) further supports this view: implicit aftereffects, intermittently probed during training with a 45° rotation, did not differ between immediate and 1.2-s delayed feedback, mirroring the present findings and reinforcing the robustness of implicit adaptation to feedback delays.

In contrast to earlier reports of progressive attenuation with increasing delays, we observed no evidence that adaptation was further suppressed across 0.2-, 0.4-, 0.8-, or 1.2-s delays. Instead, recalibration persisted at a stable, reduced level under all terminal conditions. These smaller aftereffects align with prior reports. Our larger sample (N = 50) and within-participant design provide clear evidence that the reduction reflects the use of terminal (vs. continuous) feedback, not the amount of delay: within the terminal regime (0.2–1.2 s), aftereffect magnitude did not change. Moreover, while adaptation scaled with error size under continuous feedback, this scaling was abolished across all terminal conditions, indicating that sensitivity to error magnitude depends on tight temporal coupling between movement and feedback.

By isolating effects trial by trial, our study refines models of how implicit learning depends on error type, error timing, and perturbation magnitude. The STL approach revealed subtle but reliable aftereffects that earlier block-averaging or rapid washout may have obscured, clarifying the contribution of implicit recalibration under delayed conditions. We show that early implicit adaptation reaches a saturation limit, is strengthened by task error, and requires online visual feedback. These insights extend error-based learning accounts and have practical implications for contexts where feedback is delayed or degraded, such as robotic rehabilitation, virtual training, and telemedicine. Understanding when implicit recalibration breaks down provides a basis for interventions that leverage both prediction and task errors, or that integrate multisensory and contextual cues to maintain adaptive function under suboptimal feedback.

In summary, this study shows that the limits and drivers of implicit adaptation can be revealed within a single trial. Using the STL paradigm, we found that recalibration saturates rapidly, is consistently boosted by task error, and depends on the temporal availability of feedback, though unaffected by the precise delay once feedback is terminal. These results demonstrate that saturation is intrinsic to the implicit system, that task error influences learning from the outset, and that sensitivity to error magnitude relies on real-time feedback. Beyond these theoretical insights, the STL approach itself provides a rapid, sensitive tool for mapping the dynamics of implicit adaptation, capturing effects often overlooked with block-based methods. Together, these findings refine models of motor learning and offer a foundation for translational applications aimed at enhancing recalibration in settings where feedback is delayed or error signals are ambiguous.

## Significance Statement

Motor adaptation is thought to be driven primarily by sensory prediction errors, yet the limits of this process and the influence of other error signals remain poorly understood. Using a single-trial learning paradigm, we show that implicit recalibration emerges immediately, saturates rapidly, and is consistently boosted by task error. We further demonstrate that while removing online feedback suppresses adaptation, delayed terminal feedback does not abolish it, contradicting prior claims. These findings establish saturation and task-error sensitivity as fundamental properties of the implicit system, highlight the importance of real-time feedback for error sensitivity, and introduce a rapid method for mapping adaptation dynamics. Together, this work advances mechanistic models of sensorimotor learning and informs the design of rehabilitation and training environments where feedback is delayed or degraded.

## Notes

https://osf.io/6m24e/

### Competing Interest Statement

The authors have declared no competing interest.

### Summary of Updates

Added a figure and corrected reference to figures

### Funding

- Natural Sciences and Engineering Research Council

